# Fine-tuning sequence-to-expression models on personal genome and transcriptome data

**DOI:** 10.1101/2024.09.23.614632

**Authors:** Ruchir Rastogi, Aniketh Janardhan Reddy, Ryan Chung, Nilah M. Ioannidis

## Abstract

Genomic sequence-to-expression deep learning models, which are trained to predict gene expression and other molecular phenotypes across the reference genome, have recently been shown to have poor out-of-the-box performance in predicting gene expression variation across individuals based on their personal genome sequences. Here we explore whether additional training (fine-tuning) on paired personal genome and transcriptome data improves the performance of such sequence-to-expression models. Using Enformer as a representative pretrained model, we explore various fine-tuning strategies. Our results show that fine-tuning improves cross-individual prediction performance over the baseline Enformer model for held-out individuals on genes seen during fine-tuning, with comparable performance to variant-based linear models commonly used in transcriptome-wide association studies. However, fine-tuning does not improve model generalizability on held-out genes, which contain sequences and variants unseen during fine-tuning, highlighting a remaining open challenge in the field.

## Main

Predicting the effects of genetic variation on gene expression is a key goal for advancing personalized medicine and understanding noncoding variation. Observational studies of gene expression across many individuals have identified variants in expression quantitative trait loci (eQTLs) associated with differences in the expression of particular genes. These datasets have also been used to train linear models to predict gene expression from the dosages of relatively common variants surrounding each gene (variant-based models). These models are commonly used in transcriptome-wide association studies (TWAS) [1, 2]. Recently, sequence-based deep learning models have been proposed to predict gene expression across different genes in the genome from the surrounding reference genome sequence [3–7]. Because these deep learning models use sequence rather than variant dosages as their input, they have the potential to make predictions for personal genome sequences containing any possible combination of variants. In theory, the potential advantages of sequence-based deep learning models over variant-based linear models when applied to personal genomes include the ability to (1) account for the effects of rare variants, which often have large effects on expression [8, 9]; (2) prioritize causal variant effects on gene expression rather than non-causal associations, informed by an understanding of regulatory sequence grammar learned by such models; and (3) account for non-linear effects and interactions between variants.

Such sequence-based genomic deep learning models are typically evaluated on the task of predicting gene expression and other molecular phenotypes at different held-out locations along the reference genome. They have also been evaluated on variant interpretation tasks such as distinguishing fine-mapped eQTLs from matched background variants, but such evaluations are complicated by uncertainty surrounding true causal variants in association studies, which is an ongoing challenge due to linkage disequilibrium (LD) [10]. Therefore, to directly measure performance on observed variants contained in personal genomes, two recent studies evaluated the ability of current sequence-to-expression models to explain variation in gene expression levels between individuals [11, 12]. These studies both found that current sequence-to-expression models applied to personal genome sequences are unable to explain variation in expression across individuals (cross-individual variation) for most genes. In some cases, strong negative correlations between predicted and measured expression levels suggested that models have difficulty predicting the direction of effect of regulatory variation. Additional work showed that these sequence-to-expression models also have substantial uncertainty in their predictions on personal genomes [13].

Because existing sequence-to-expression models were trained on sequences solely from the reference genome, we hypothesized that seeing examples of genetic variation and paired molecular phenotype data during training would enable such models to better learn the effects of small sequence differences. To test this hypothesis, we perform a series of evaluations to determine whether fine-tuning an existing sequence-based genomic deep learning model on paired personal genome and transcriptome data in lymphoblastoid cell lines (LCLs) from the Geuvadis consortium [14] improves performance on cross-individual prediction of gene expression from personal genomes. We use Enformer [6] as a pre-trained genomic deep learning model that has been widely used to model gene expression, chromatin accessibility, histone modifications, and transcription factor binding across many cell types. We test multiple strategies for fine-tuning the pre-trained Enformer model and compare performance relative to the baseline (non-fine-tuned) Enformer and to several variant-based models commonly used in the TWAS field. We use an input context size of *∼*49.2 kb for training and evaluating all models. Our fine-tuning strategies (see Methods for details) include single sample regression (in which each individual’s personal expression level and paired personal sequence for each gene is treated as an independent training example), pairwise regression and pairwise classification (in which the predicted and measured expression levels for pairs of individuals are compared), and joint training on both the personal data and either Enformer’s original training data or massively parallel reporter assay (MPRA) data. A schematic of the pairwise regression fine-tuning strategy, which is used throughout the main figures, is shown in Fig. 1. See Fig. S1 for schematics of other fine-tuning strategies.

**Fig. 1.**
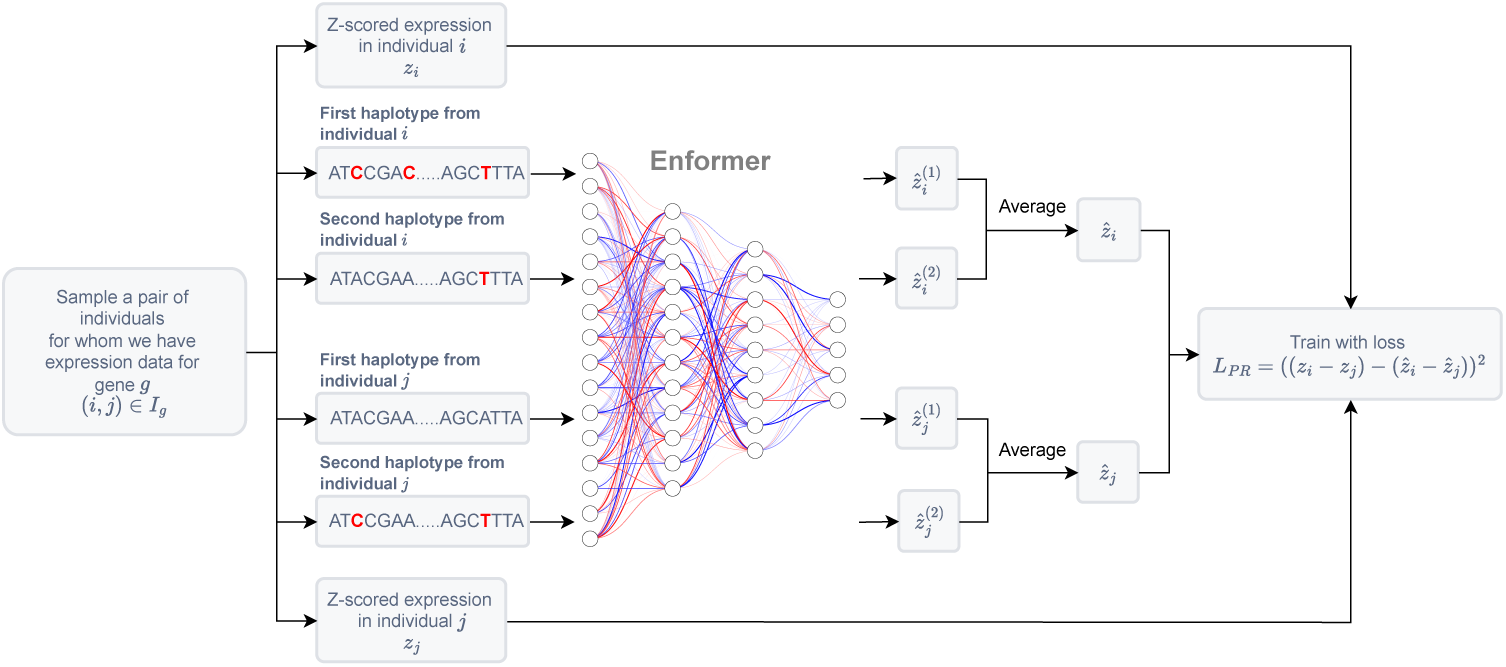
Schematic of the pairwise regression method used to fine-tune Enformer on paired personal genome and transcriptome data. For fine-tuning, a gene *g* is sampled from a set of genes *G*. From the training individuals *I_g_* for gene *g*, a pair of individuals (*i, j*) is randomly sampled, along with their z-scored expression levels *z_i_* and *z_j_*. Enformer is fine-tuned to predict the expression difference *z_i_ − z_j_* from the individuals’ personal genome sequences using a mean squared error loss between the actual difference and the predicted difference *ẑ_i_ − ẑ_j_*. Red nucleotides highlight genetic variants in each haplotype.

We evaluate our fine-tuned models in three main settings of increasing difficulty (Fig. 2a). First, we assess whether fine-tuning with personal data from 200 genes enables the models to generalize to randomly-held out individuals for the same genes (“random-split” genes). This mirrors the standard evaluation of variant-based linear models used in TWAS. Second, variant-based models have been shown to have notably reduced performance when applied to individuals from populations different from those in the training set [15, 16]. Therefore, we evaluate whether seeing personal data from only European ancestry individuals during fine-tuning for another set of 200 genes results in models that generalize to Yoruban ancestry individuals for the same genes (“population-split” genes). Because the fine-tuned models are initialized with weights from Enformer, which should have learned some regulatory sequence grammar during pre-training, these models might be more likely than variant-based models to attribute regulatory effects to causal variants rather than non-causal associations within strong LD blocks. Third, we evaluate performance of the fine-tuned models on genes for which no personal data was seen during fine-tuning (“unseen” genes).

**Fig. 2.**
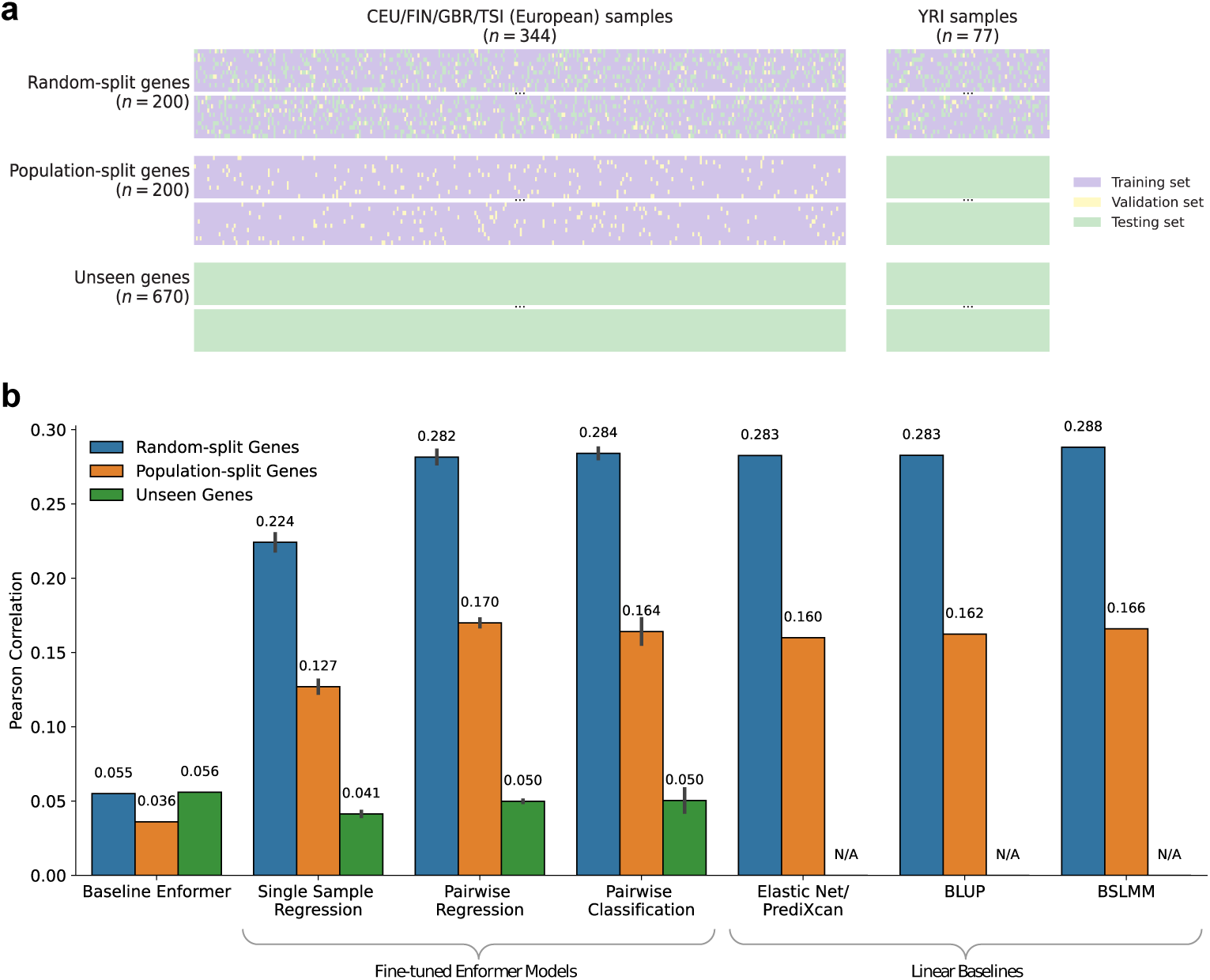
**(a)** The three gene sets used for model evaluation: random-split genes, population-split genes, and unseen genes. Each row represents a gene and each column an individual, with colors indicating membership in the training, validation, or testing sets. 20 example genes are shown for each gene set. Individuals in our dataset come from five populations: CEU (Utah residents with Northern and Western European ancestry), FIN (Finnish in Finland), GBR (British in England and Scotland), TSI (Toscani in Italy), and YRI (Yoruba in Ibadan, Nigeria). **(b)** Performance of fine-tuned Enformer models compared to baseline methods in predicting gene expression across individuals. We trained three replicates for each of the fine-tuned Enformer models using different random seeds, which shuffled the data order and pairs sampled (for pairwise models) while keeping the set of training individuals per gene fixed. Bar heights indicate the mean (over genes) of the cross-individual Pearson correlation between measured and predicted expression levels. For fine-tuned models, bar heights are averaged across three replicates, with error bars showing the standard deviation.

In random-split genes, held-out individuals have input sequences that do not exactly match any of the sequences seen by models during training or fine-tuning, so this evaluation tests model generalizability to new sequences with different combinations of variants than in the training set. However, many specific variants present in randomly held-out individuals—particularly common variants—are still the same as those found in the training set individuals (Fig. S2). On this task (Figs. 2b and S3, blue bars, and Fig. S4a), we find a mean test set Pearson correlation between predicted and measured expression levels of just over 0.28 for most of the fine-tuned models. This is a substantial improvement over the baseline Enformer model (0.055) and is comparable to variant-based models trained for the same set of genes using the same training samples. This improvement over baseline Enformer indicates that fine-tuning on personal genome and transcriptome data enables sequence-based models to more accurately learn variant effects on expression, or at least variant associations with expression, including correcting the direction of effect errors previously observed. We note that several of the fine-tuning strategies and several variant-based models all reach very similar levels of performance, suggesting that they all learn similar information from the personal data.

On population-split genes for which we hold out Yoruban individuals (Figs. 2b and S3, orange bars, and Fig. S4b), we observe a drop in performance relative to the evaluation on random-spit genes for both the variant-based linear models and the finetuned sequence-based models. This drop is consistent with previous observations for variant-based models and, in part, reflects the larger fraction of variants in the held-out population that are unobserved in the training individuals (Fig. S2). This drop also reflects the fact that some variant effects learned by the models may actually be non-causal associations present in the training data, which are consistently correlated with causal variants in individuals with the same LD structure, but are less predictive in populations with different LD structure where the relevant correlation is reduced. The similar drop in performance that we observe for the fine-tuned models and the variant-based models suggests that all tested models are prone to learning non-causal associations. The top-performing fine-tuned models (e.g. the pairwise regression model, with Pearson correlation 0.17) slightly out-perform the variant-based models, which could reflect a slight improvement in learning effects for causal variants rather than correlated non-causal variants.

We perform two additional evaluations to more directly test whether the fine-tuned models are better able to identify causal variants. First, we consider a set of recently fine-mapped LCL eQTLs, which should be enriched for causal variants, from a large and globally-diverse study (Multi-ancestry Analysis of Gene Expression, MAGE) [17], restricted to the genes and common variants seen during fine-tuning. For the pairwise regression fine-tuned model, the linear variant-based models, and baseline Enformer, we determine the fraction of fine-mapped variants that are predicted to have high effect sizes (Fig. 3a). We find that the fine-tuned model ranks these putatively causal fine-mapped variants more highly than the variant-based models and baseline Enformer, suggesting that fine-tuned deep learning models are more likely to identify true causal variants. Next, we consider a set of expression-altering variants identified in an MPRA study in the same cell type [18], again restricted to the genes and common variants seen during fine-tuning, and perform a similar analysis (Fig. 3b). We find that the fine-tuned model and the baseline Enformer model both rank these MPRA-effect variants more highly than the variant-based models. In this case, the baseline Enformer model outperforms the fine-tuned model, indicating that the variant effects learned from personal variation in an endogenous context during fine-tuning are not entirely transferable to the MPRA context. This result, in conjunction with those above, suggests that while the pre-trained Enformer model understands regulatory principles relevant in the MPRA context, the effects of expression regulation are different in the local context of an MPRA than in the endogenous genome, consistent with previous comparisons of MPRA and eQTL studies [19].

**Fig. 3.**
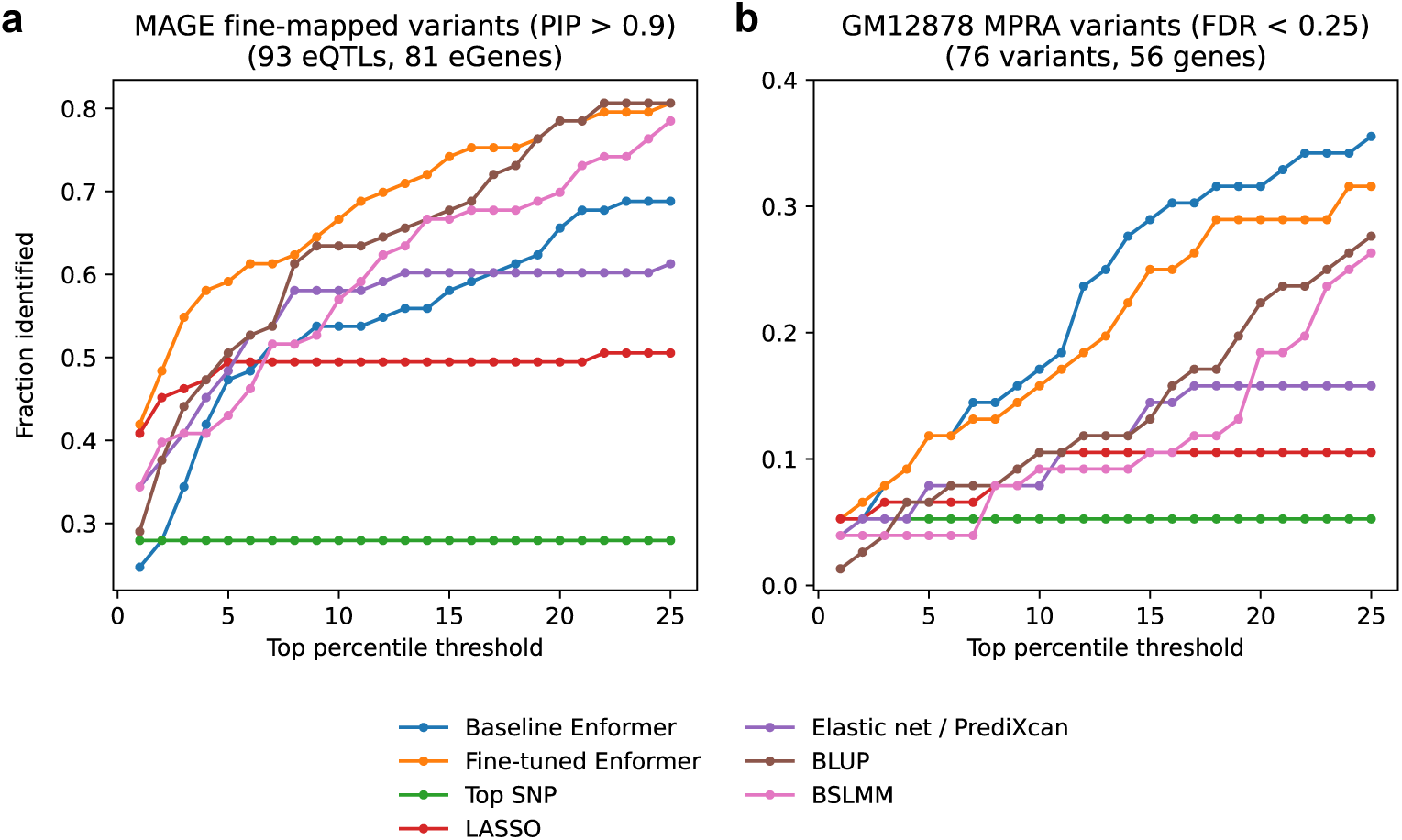
Model performance in identifying variants likely influencing gene expression in LCLs. We evaluated the ability of models to prioritize putatively causal common variants within the *∼*49.2 kb context window around random-split and population-split genes. These variants come from two sources: **(a)** fine-mapped variants from the MAGE eQTL study [17] with a poster inclusion probability (PIP) *>* 0.9 and **(b)** variants from a GM12878 MPRA [18] that significantly influence promoter expression with a false discovery rate (FDR) *<* 0.25. For each model, we calculated the percentile rank of each putatively causal variant’s predicted effect magnitude (*|β|* for linear models and *|*ISM*|* for Enformer models) relative to the predicted effect magnitudes for all other common variants within the context window of the same gene. We plot the fraction of putatively causal variants identified (y-axis) across different top percentile thresholds (x-axis).

On unseen genes, model performance reflects generalizability to unseen sequences containing unseen variants, which tests whether seeing paired genome and expression variation during fine-tuning has improved the cis-regulatory grammar learned by the models in a way that better captures causal variant effects. In addition, a model that can generalize to unseen genes should be similarly able to generalize to unseen variants in seen genes, including rare or *de novo* variants. Variant-based models cannot be applied to such variants, since they learn weights only for variants that are observed at high enough frequency during training. Therefore, for this evaluation, we compare performance of only the fine-tuned models and baseline Enformer. Despite the improved performance of fine-tuned models over baseline Enformer for held-out individuals in seen genes, we find that their performance on unseen genes is comparable to or slightly worse than baseline Enformer (Figs. 2b and S3, green bars). This result also holds when considering the absolute value of the cross-individual correlations for fine-tuned and baseline Enformer models (Fig. S5). Thus the fine-tuned models have not learned a more generalizable representation of variant effects or their magnitudes. The performance of the fine-tuned models on different individual unseen genes varies widely, including many genes with negative correlations between measured and predicted expression levels (Fig. S6), as was previously observed for baseline Enformer [11, 12]. Comparing the performance of the pairwise regression fine-tuned model and baseline Enformer on individual unseen genes (Fig. S6), we also find a similar pattern as previously observed when comparing different deep learning models or model replicates [11, 13], with greater consistency in the magnitude than the direction of correlation.

The similar performance of the fine-tuned models and the variant-based linear models on held-out individuals suggests that the fine-tuned models are primarily learning linear combinations of the effects of common variants, rather than non-linear effects or interactions between variants. Indeed, when we construct a linear approximation of the pairwise regression fine-tuned model using its *in silico* mutagenesis (ISM) scores as the weights for variant dosages in a linear model, we find very similar performance between the full fine-tuned model and the linear approximation (Fig. 4). This analysis indicates that although deep learning models have the capacity to learn complex interactions, their predictions of variant effects from personal genomes are primarily linear without strong interactions between variants. Note that this analysis does not mean that genomic deep learning models do not learn any interactions between positions within an input sequence (such as interactions between neighboring bases within transcription factor binding motifs), but simply that the learned effects of variants at their typical spacing found within personal genomes (*∼*1kb) are approximately linear and additive.

**Fig. 4.**
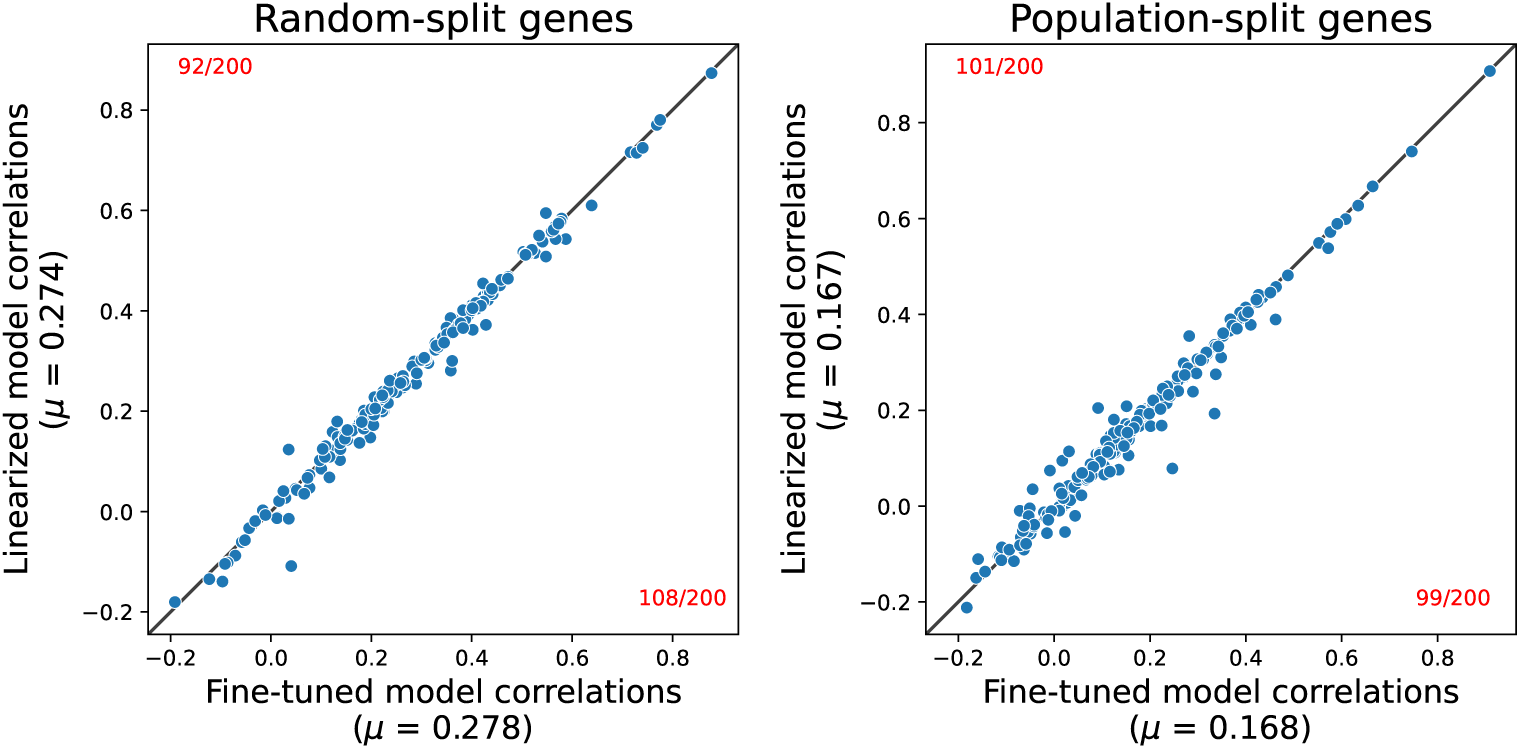
Performance comparison of the full fine-tuned Enformer model (pairwise regression) to a linear approximation, termed the linearized model. For random-split genes (left) and population-split genes (right), pairwise scatterplots display the cross-individual Pearson correlations of the two models, with each point representing a distinct gene. A black *y* = *x* line serves as a reference, and the number of genes lying above and below this line is highlighted in red. The linearized Enformer model is constructed as a weighted linear combination of variant dosages, where the weights are *|*ISM*|* scores.

To determine how much the predictions of the fine-tuned sequence-based models depend on common versus rare variants, we also evaluate the pairwise regression fine-tuned model on partial personal genome sequences, in which we only include personal variants whose minor allele frequency (MAF) exceeds a threshold. By varying this threshold (Fig. 5), we find that common variants contribute the most to cross-individual performance, but that there is still a small contribution from variants that were rare (*<*5% MAF) in the training set individuals. We also investigate the individual variants that contribute most to the model predictions (drivers), and find similar numbers of drivers for the pairwise regression fine-tuned model and the baseline Enformer model (Fig. S7). Drivers for both models include a number of rare variants (*<*5% MAF) that are not captured in the variant-based models, though the fine-tuned Enformer drivers are on average more common than those of baseline Enformer. In addition, baseline Enformer drivers are predominantly located close to the gene start site (within *∼*5kb), while the variant-based model drivers are roughly uniformly distributed within the input context window, and the fine-tuned model drivers have an intermediate distribution.

**Fig. 5.**
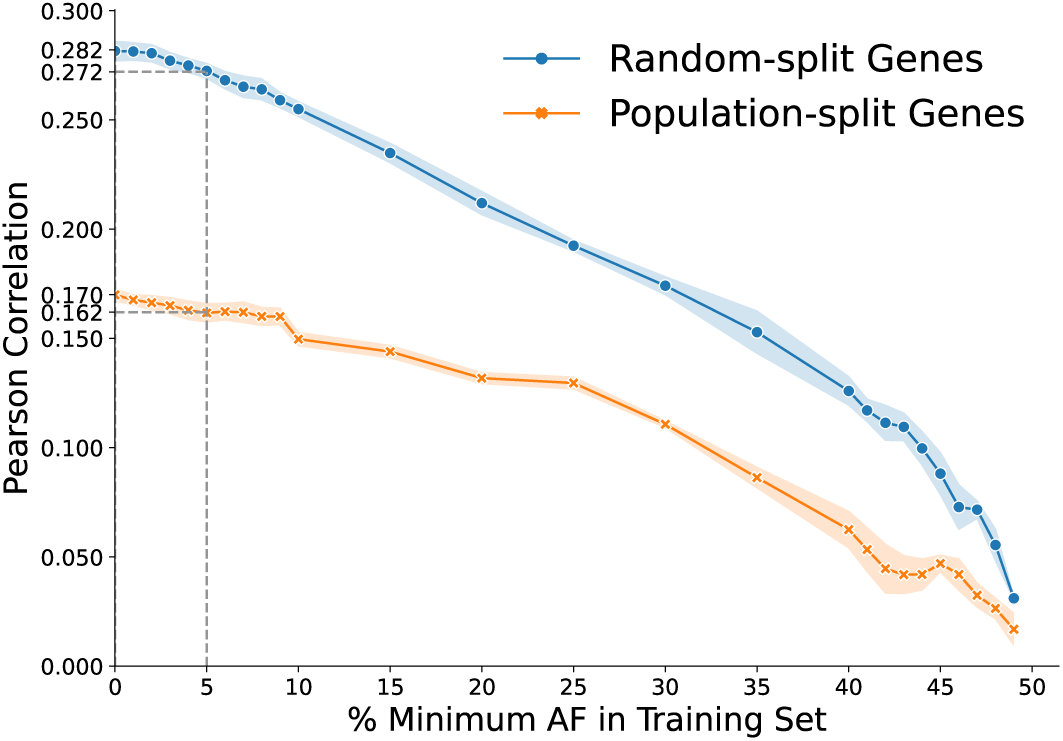
Contribution of rare variants to the performance of the fine-tuned Enformer model (pairwise regression). Minor alleles with an allele frequency (AF) in training samples below the threshold shown on the x-axis were replaced with the corresponding major allele to construct partial personal genome sequences, which were then input into the model to generate predictions. The y-axis displays the mean (over genes) of the cross-individual Pearson correlation, averaged over three replicates, with shaded regions showing the standard deviation. Gray dashed lines indicate the performance on random-split and population-split genes when rare variants (AF *<* 5%) are excluded.

In conclusion, we find that multiple strategies for fine-tuning the Enformer sequence-to-expression model using observed genome and transcriptome variation improve performance on predicting cross-individual variation in gene expression from personal genomes, a task with which the original Enformer model struggles. In agreement with recent results from a similar fine-tuning study [20], we find that this improved performance is comparable to that of variant-based linear models trained on the same genes. Our results indicate that variant-based approaches, which explicitly use variant dosages as predictors, and sequence-based approaches, which model the full sequence context, are similarly able to learn variant associations with gene expression for the specific genetic variants observed in their training data. Both approaches also have similar scaling with the number of individuals seen during training, but appear to be reaching a plateau (Fig. 6), suggesting that training on even larger numbers of individuals may not substantially improve performance, likely due to the large amount of shared variation between individuals in human populations. Alternative approaches to increase the amount of sequence diversity seen during training may be needed, although fine-tuning jointly with MPRA data did not improve performance in our analyses. In addition, although fine-tuning on personal data enables the sequence-based models to learn observed associations between variants and gene expression levels, the fine-tuned models do not generalize to new sequences with new variants, as reflected in their poor performance on unseen genes. Developing sequence-based models with an understanding of variant effects on regulatory sequence grammar that is fully generalizable to unseen sequences and variants will likely require new types of data or new strategies for utilizing observed variation during training, highlighting an important area for future work in the field.

**Fig. 6.**
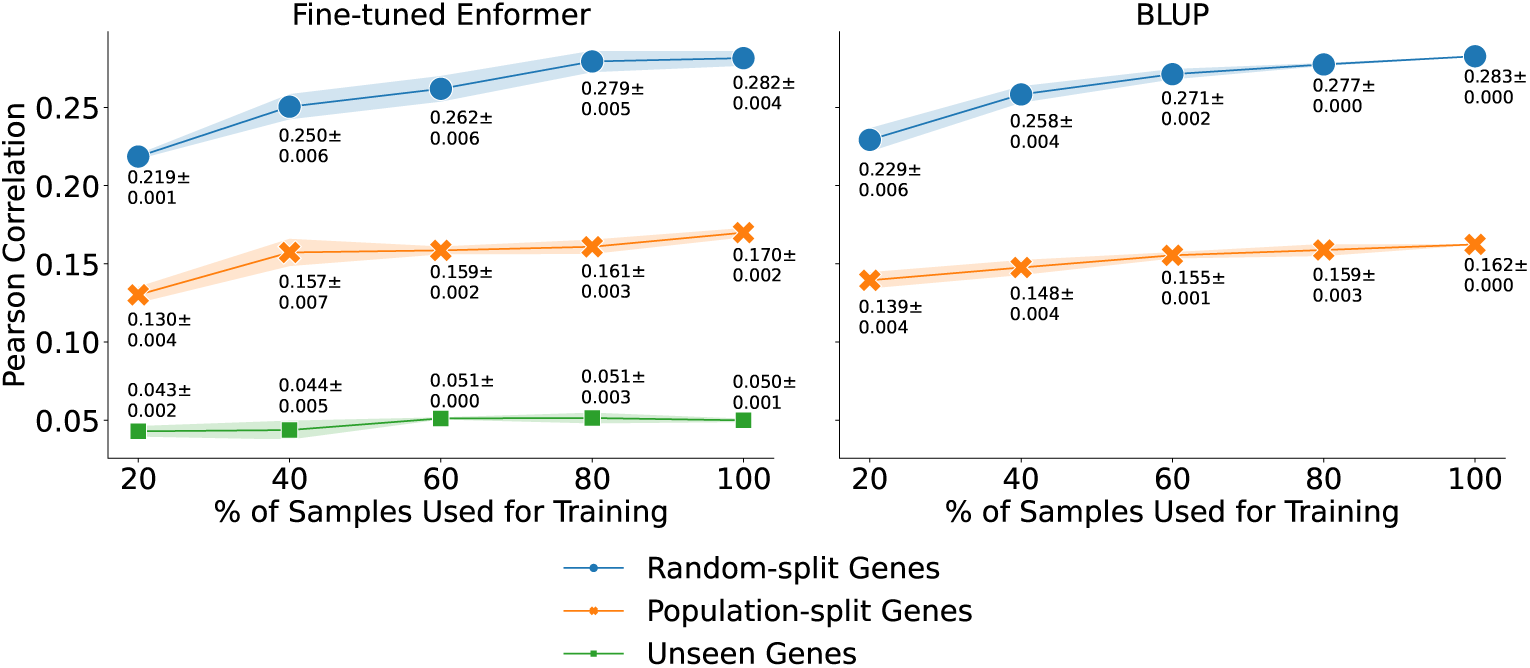
Performance comparison of fine-tuned Enformer (pairwise regression, left) and BLUP (right) as a function of the fraction of training individuals used. Mean cross-individual Pearson correlations and standard deviations are calculated across three replicates, each trained on a different subsampled set of individuals. BLUP cannot make predictions for unseen genes; therefore, performance metrics for unseen genes are excluded from the BLUP panel.

## Methods

### Gene expression dataset

We trained and evaluated models using RNA sequencing (RNA-seq) data from lymphoblastoid cell lines (LCLs) provided by the Geuvadis consortium [14]. This dataset includes 421 individuals with corresponding phased whole genome sequencing (WGS) data from the 1000 Genomes Project [21]. These 421 samples come from five populations: 92 Tuscan (TSI), 89 Finnish (FIN), 85 British (GBR), 78 European individuals from Utah (CEU), and 77 Yoruban (YRI).

To prepare the RNA-seq data, we preprocessed the uncorrected RPKM data from the `GD660.GeneQuantRPKM.txt.gz` file. First, to facilitate better comparisons of RNA abundances across samples, we converted gene expression measurements from RPKM to TPM using the formula

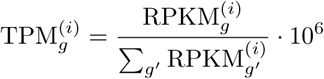

where *g* denotes a specific gene and *i* denotes a specific individual. We then excluded poorly-expressed genes with zero read counts in at last 50% of the samples. To make the expression distribution of each gene closer to a Gaussian, we applied a log transformation to the TPM values, using a pseudocount of 0.01. Finally, to correct for potential batch effects and account for population structure, we regressed out the first 10 principal components [22].

### Data splits

To assess model performance, we constructed three disjoint datasets of increasing difficulty. From the 3,259 genes with a statistically significant eQTL (FDR *<* 0.05) in the Geuvadis EUR *cis*-eQTL analysis, we randomly selected (1) 200 genes for random-split evaluation, designed to test model performance on held-out individuals; (2) 200 genes for population-split evaluation, intended to assess generalization across populations; and (3) 670 genes for unseen-gene evaluation, aimed at evaluating model performance on entirely new genes and variants. To prevent data leakage, we ensured that genes selected for each evaluation set were located on different chromosmes.

For random-split genes, we randomly partitioned the 421 individuals into a development set of 344 samples and a testing set of 77 samples, irrespective of their population. The development set was further divided into a training set of 327 individuals and a validation set of 17 individuals. For population-split genes, all 344 individuals from the European super-population (comprising TSI, FIN, GBR, and CEU populations) formed the development set, which was further randomly divided into training and validation sets of the same size as before. The testing set consisted of the 77 Yoruban (YRI) individuals. For unseen genes, all 421 individuals were in the testing set.

### Fine-tuning

We fine-tuned Enformer [6] to predict gene expression in LCLs from personalized genome sequences. As in Huang et al. [11], personalized genome sequences for each individual’s haplotype were generated using `bcftools consensus` [23] with hg19 as the reference genome, consistent with the Geuvadis variant calls. Only single-nucleotide variants were incorporated into these sequences. For computational tractability, we shortened personal genome sequences to span the 49,152 base pairs centered around the transcription start site (TSS) of each gene, as annotated in Gencode.v12—which served as the same reference for RNA-seq quantification. This sequence length corresponds to one-fourth of the maximum input length that Enformer can accept.

We fine-tuned the entire Enformer model–without freezing any weights–using multiple approaches detailed below. All models were trained jointly on the 400 combined random-split and population-split genes. During training, we augmented the dataset by randomly reverse-complementing input sequences and shifting them by up to three base pairs in either direction (padding with zeros as necessary). We fine-tuned using the AdamW optimizer [24] with a learning rate that linearly increased to its peak over 1,000 steps and then decayed to zero using cosine annealing over 50 epochs. Experiments were conducted using eight GPUs (NVIDIA A5000s or TITAN RTXs) with a batch size of eight for most models and sixteen for the single sample regression model.

Two hyperparameter configurations were evaluated: (1) a learning rate of 5 *×* 10*^−^*^4^ and weight decay of 5 *×* 10*^−^*^3^, and (2) a learning rate of 1 *×* 10*^−^*^4^ and weight decay of 1 *×* 10*^−^*^3^. The configuration yielding the lowest validation loss was selected. To prevent overfitting, early stopping with a patience of five epochs based on the validation loss was implemented. During inference, predictions were obtained by averaging the four predictions from the forward and reverse strand sequences of both haplotypes.

We now describe the specific model architectures and loss functions used in our various approaches.

#### Single sample regression

The single sample regression model directly predicts a single individual’s gene expression from their two haplotypes (Fig. S1a). Because Enformer does not natively have an LCL RNA-seq output track, we introduced an additional output head to generate predictions. Briefly, Enformer (using convolution and transformer layers) generates embeddings of size 384 *×* 3072 for each haplotype.

The central ten bins of the embedding are isolated and aggregated using attention pooling, which performs a weighted linear combination. The pooled embeddings are then projected into predictions via a linear layer. The final gene expression prediction is obtained by averaging the predictions from both haplotypes.

Because a naive mean-squared error (MSE) loss would bias the model towards genes with high variance, we employ the symmetric mean absolute percentage error (SMAPE) loss to quantify the relative error between the predicted value *ŷ* and the true value *y*:

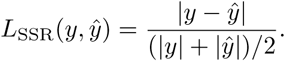

We found that a learning rate of 1*×*10*^−^*^4^ and weight decay of 1*×*10*^−^*^3^ was the optimal hyperparameter configuration.

#### Pairwise regression

To emphasize genomic positions where individuals differ, we adopted a pairwise regression strategy inspired by contrastive learning (Fig. S1b). In this approach, we input the haplotypes of two individuals into the model. The model shares the same architecture as the single sample regression model to generate predictions for each individual. However, instead of penalizing individual prediction errors, the model is trained to predict the expression difference between two individuals using the following pairwise loss:

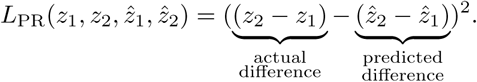

We utilized z-scored expression values to standardize gene variances, ensuring that all genes contribute equally to the loss. We found that a learning rate of 1 *×* 10*^−^*^4^ and weight decay of 1 *×* 10*^−^*^3^ was the optimal hyperparameter configuration.

#### Pairwise classification

In previous approaches, we introduced a new output head to make RNA-seq predictions. However, adding parameters could enable the model to adapt by only updating new components without updating the core parameters essential to learn a generalizable cis-regulatory grammar. Therefore, in this approach, we leveraged the model’s existing predictions for the CAGE track of a corresponding lymphoblastoid cell line, GM12878.

Because the CAGE outputs differ significantly in scale from the RNA-seq data used for fine-tuning, we reformulate the task as a binary classification problem (Fig. S1c). Specifically, we train the model to predict whether the gene expression of one individual exceeds that of another. First, we sum the ten central bins of the GM12878 CAGE track for each haplotype to obtain a single expression prediction per individual. The predictions for individuals 1 and 2, *ŷ*_1_ and *ŷ*_2_, follow a Poisson distribution, since Enformer outputs are Poisson. Computing *P* (*ŷ*_1_ *> ŷ*_2_) requires computing the cumulative distribution function of a Skellum random variable, which is intractable. To address this, we apply the Anscombe transform to approximate the Poisson distributions as normal distributions, making the above probability computation tractable. From this, we calculated the binary cross-entropy loss between the predicted probability and the ground truth:

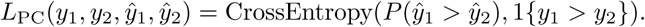

To reduce the effects of measurement noise, we applied the classification loss only to pairs whose expression percentiles for the given gene differed by at least 25%. We found that a learning rate of 1 *×* 10*^−^*^4^ and weight decay of 1 *×* 10*^−^*^3^ was the optimal hyperparameter configuration.

#### Joint training with Enformer’s original dataset

In prior approaches, we finetuned Enformer exclusively on personalized expression data. However, neural networks are prone to catastrophic forgetting, a phenomenon where training on new tasks can cause models to lose information learned in previous tasks [25, 26]. In our case, this could result in Enformer forgetting the regulatory grammar learned during its original training, resulting in poor performance especially on unseen genes.

To mitigate this [27, 28], we jointly trained Enformer on both the personalized expression dataset and its original training dataset. Enformer’s original dataset consists of 196,608 base pair sequences from human and mouse reference genomes, along with associated gene expression, chromatin accessibility, histone modification, and transcription factor binding measurements (5,313 human and 1,643 mouse). We utilized the same sequences from the original Enformer dataset as were used by Avsec et al. [6] for training. During joint training, each batch included an equal number of sequences from the original human dataset, original mouse dataset, and personalized expression dataset. We employed Enformer’s standard architecture and Poisson loss for the original data, while applying either our pairwise regression and pairwise classification architectures and loss functions to the personalized expression data. We found that a learning rate of 5 *×* 10*^−^*^4^ and weight decay of 5 *×* 10*^−^*^3^ was the optimal hyperparameter configuration.

#### Joint training with MPRA data

To augment the genetic variation seen by Enformer beyond that present in our personal genomes dataset, we fine-tuned Enformer using both personalized expression data and Massively Parallel Reporter Assay (MPRA) data. MPRAs are high-throughput assays that assess the impact of short regulatory sequences, used as promoters, on downstream reporter gene expression in host cells, separate from their endogenous context. MPRAs can quantify variant effects by measuring the log-fold change in expression induced by promoter segments containing a variant compared to their corresponding reference segments. We utilized the MPRA dataset from Siraj et al. [18], which quantifies the effects of over 200,000 trait-associated variants on promoter-driven expression across five cell types: GM12878, HepG2, K562, A549, and SK-N-SH (see Table S1 for ENCODE [29–31] accession IDs).

We fine-tuned on the MPRA data using a pairwise regression approach, analogous to our previous approach for personal expression data. For each variant, both the 200 bp reference and alternate sequences were input into the model. Cell-type specific expression predictions were generated for each sequence using separate linear output heads that processed the flattened Enformer embeddings. The predicted cell-type specific effect was calculated as the difference between alternate and reference expression predictions. This prediction was compared to the z-scored ground-truth effect size from the MRPA dataset using the loss function

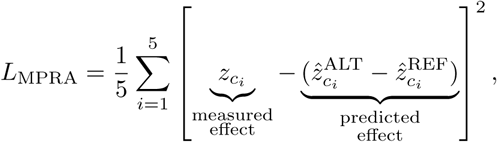

where *c_i_* denotes the *i*-th cell type. If a variant effect was unavailable for a particular cell type, the corresponding loss term was excluded.

During joint fine-tuning, each batch contained an equal number of examples from the MPRA dataset and the personalized expression dataset. We found that a learning rate of 1 *×* 10*^−^*^4^ and weight decay of 1 *×* 10*^−^*^3^ was the optimal hyperparameter configuration.

### Baselines

#### Baseline Enformer

We generated predictions using the original Enformer model on the same personal genome sequences that were used for fine-tuning, which included only SNPs within the approximately 49.2 kb region centered at the gene’s TSS. For each input sequence, we obtained a prediction by averaging the center 10 output bins of the GM12878 CAGE track. Individual-level predictions were then calculated by averaging the four predictions from the forward and reverse strand sequences of both haplotypes.

#### Variant-based linear methods

We also benchmarked five variant-based linear methods commonly used in TWAS: top SNP, LASSO [32], elastic net (also known as PrediXcan) [1, 33], BLUP [34], and BSLMM [35]. These methods, implemented using the FUSION package [2], produce a separate model for each gene. Training and testing were conducted on both random-split and population-split genes, using the same sample splits as in fine-tuning. All models were trained to predict z-scored expression values from the standardized dosages of common SNPs (MAF *≥* 5% in training set samples) within the approximately 49.2 kb context window centered at the gene’s TSS. We did not observe a performance improvement when the models were allowed to use variants within the full 1 Mb context window commonly recommended (data not shown). The optimal hyperparameter *λ* in the elastic net model was chosen through 5-fold cross-validation on the training set, consistent with FUSION’s default settings. The other methods did not have tunable hyperparameters.

### Computing *in silico* mutagenesis (ISM) scores

For the Enformer models, we calculated ISM scores for all variants in our dataset. The ISM score for each variant is determined by the formula ISM = *f* (*x*_ALT_) *− f* (*x*_REF_), where *f* predicts gene expression in LCLs. Here, *x*_REF_ represents the reference hg19 sequence centered at the gene’s TSS, and *x*_ALT_ is the same sequence with the variant’s alternate allele substituted at the appropriate position. We compute ISM scores separately for the forward and reverse strand sequences and then average these two values to obtain the overall ISM score. Similar to coefficients (*β*’s) in a linear model, ISM scores reflect the marginal change in gene expression when the dosage of the variant increases by 1.

### Identifying putatively causal variants

We evaluated the ability of models to identify individual variants that have been previously associated with likely changes in gene expression in LCLs. We obtained putatively causal variants from two sources.

First, we selected fine-mapped eQTLs with a posterior inclusion probability (PIP) *>* 0.9 according to SuSiE [36] from the MAGE eQTL study (DOI: 10.5281/zenodo.10535719) [17]. We limited this set of fine-mapped eQTLs to those used by all models in training by applying the following pipeline. Initially, we mapped eQTLs from hg38 to hg19 using `Picard LiftoverVcf` [37]. Then, we retained eQTLs within the context window of one our training genes (random-split and population-split) that were present in our training set with a minor allele frequency (MAF) *≥* 5%. We used fine-mapped eQTLs from MAGE instead of Geuvadis because the MAGE dataset is larger (731 versus 462 individuals) and has greater genomic diversity.

Second, we obtained variants that significantly alter promoter expression (FDR *<* 0.25 after Benjamini-Hochberg correction) from a GM12878 MPRA [18] (same as used for joint fine-tuning with MPRA data), whose variant effect quantifications can be found at the ENCODE accessions listed in Table S1. We again retained only those variants within the context window of one of our training genes and with an MAF *≥* 5% within our training set.

For each model, we calculated the percentile of a putatively causal variant’s absolute effect size relative to the absolute effect size of all other variants within the context window of the gene that have a training MAF *≥* 5%. The absolute effect size is *|β|* for linear models and *|*ISM*|* for Enformer models. When multiple variants had identical absolute effect sizes, the percentile was determined by averaging their ranks.

### Linear approximation to Enformer

For each gene *g*, we constructed a linear function 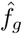 that uses a weighted combination of variant dosages *x*_1_*,…, x_n_* (with ISM scores as weights) to approximate the predictions of the non-linear Enformer model:

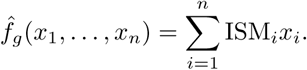

This approximation is not exact if the original model contains non-linear dosage effects or interaction terms between variants.

### Rare variant analysis

We assessed the impact of rare variants on the fine-tuned Enformer model’s predictions by replacing minor alleles with major alleles in personal genome sequences during inference. This substitution occurred when allele frequencies—calculated using the training set for each gene—fell below specified thresholds. By applying this procedure across various frequency thresholds, we evaluated the contribution of rare variants to the model’s predictions.

### Driver analysis

To identify significant genetic variants (drivers) that influence model predictions for each gene, we employed an algorithm that accounts for correlations between variants, first proposed in Sasse et al. [12]. We start by approximating the model’s potentially non-linear prediction function with a simpler linear function. For variant-based linear models, this approximation is exact; for Enformer models, we use the linear approximation discussed above (the accuracy of this approximation was validated in Fig. 4).

We rank all variants based on the magnitude of their effect sizes—coefficients for linear models and ISM scores for Enformer models. Starting with the most significant variants, we iteratively add a variant to the driver set if it meets two criteria. First, including the variant must increase the Pearson correlation between our linear approximation and the original model’s predictions by at least 0.05. Second, the variant’s individual contribution must have a significant (Bonferroni-corrected p-value *<* 0.01 and positive correlation with the model’s predictions.

## Code and data availability

Code to reproduce our analyses and to fine-tune Enformer is available at https://github.com/ni-lab/finetuning-enformer. The Geuvadis gene expression data and whole genome sequencing data used in this study are publicly available at https://www.ebi.ac.uk/biostudies/arrayexpress/studies/E-GEUV-1. Enformer’s original training data were derived from the data published at https://console.cloud.google.com/storage/browser/basenji_barnyard/data. MPRA data were downloaded from ENCODE using the accession IDs listed in Table S1.

## Acknowledgements

We thank Sara Mostafavi, Maria Chikina, Katherine Pollard, Shiron Drusinsky, Sean Whalen, Cory McLean and members of the Ioannidis lab for helpful discussions. This work was partially supported by the Berkeley Artificial Intelligence Research (BAIR) Open Research Commons, and used the Savio computational cluster resource provided by the Berkeley Research Computing program at the University of California, Berkeley (supported by the UC Berkeley Chancellor, Vice Chancellor for Research, and Chief Information Officer). N.M.I. is a Chan Zuckerberg Biohub San Francisco Investigator.

## Supplementary Figures

**Fig. S1.**
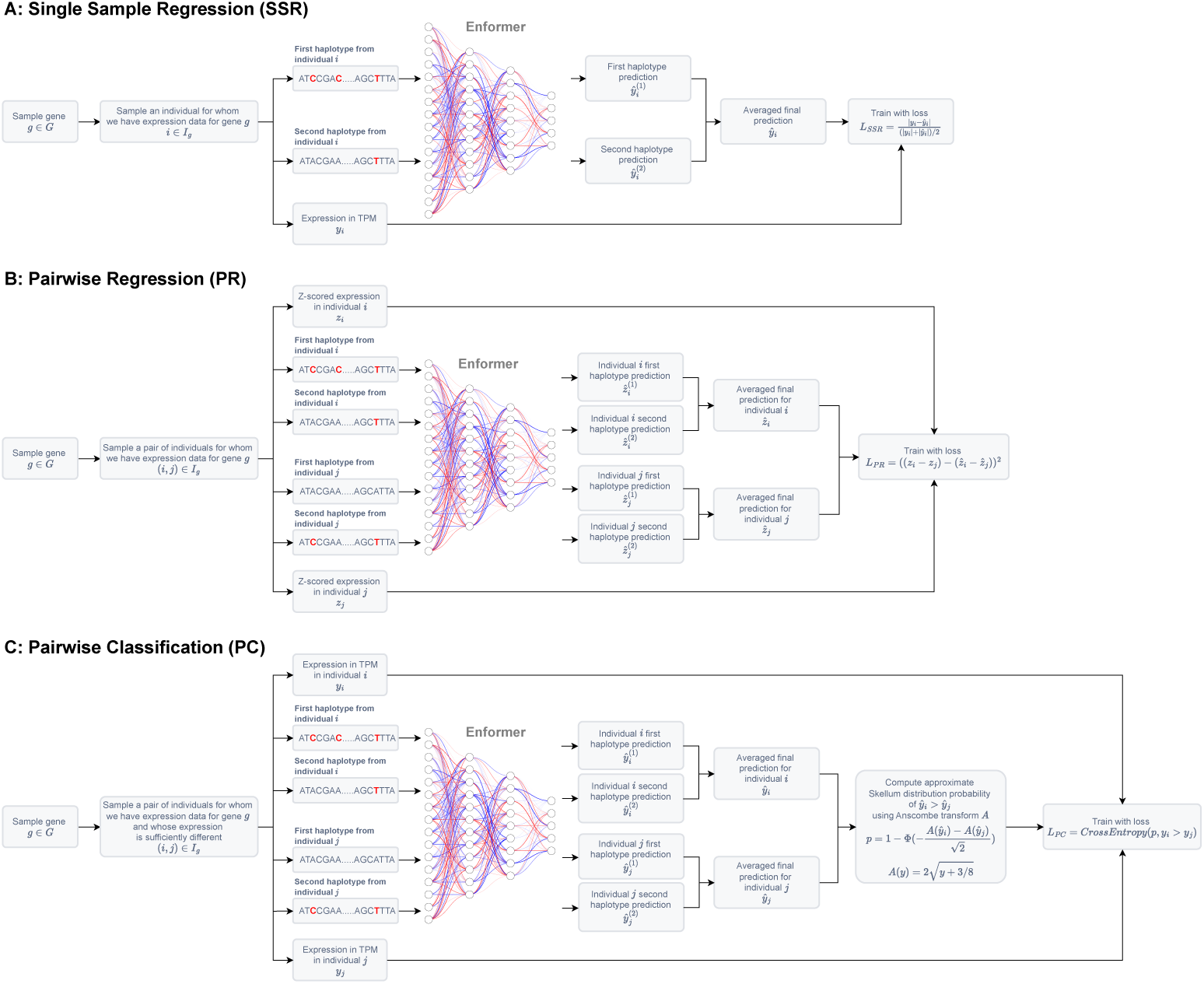
Schematics of the three primary methods used to fine-tune Enformer on paired personal genome and transcriptome data: **(a)** single sample regression, **(b)** pairwise regression, and **(c)** pairwise classification.

**Fig. S2.**
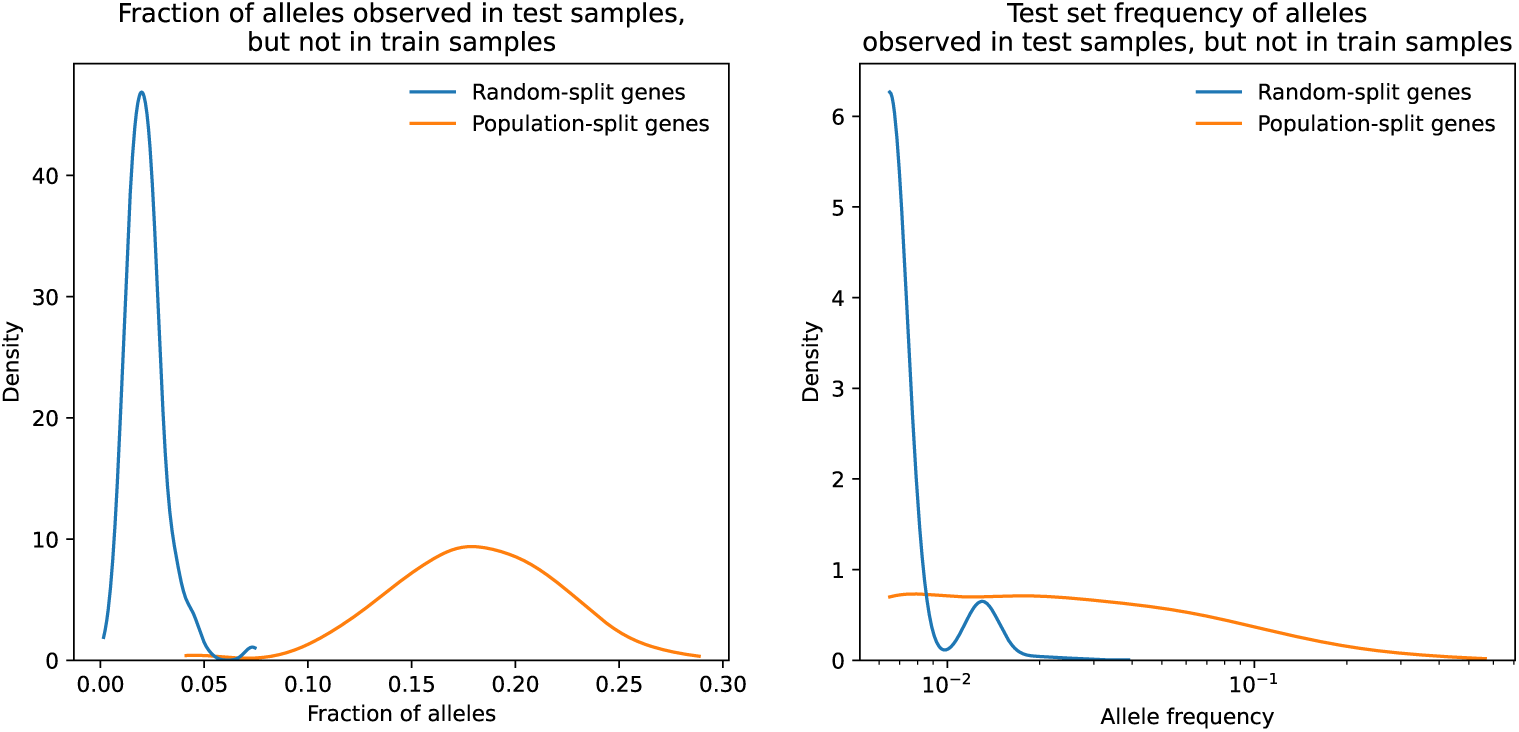
**Left**: For each gene, we computed the fraction of alleles at polymorphic (non-fixed) positions observed at least once in the test set but absent in the training set. A kernel density estimate (KDE) visualizes the distribution of these fractions, stratified by gene set. **Right**: The allele frequency distribution of alleles observed in the test set but absent in the training set at polymorphic positions is shown, using a KDE. Frequencies are computed with respect to individuals in the test set, with data similarly stratified by gene set.

**Fig. S3.**
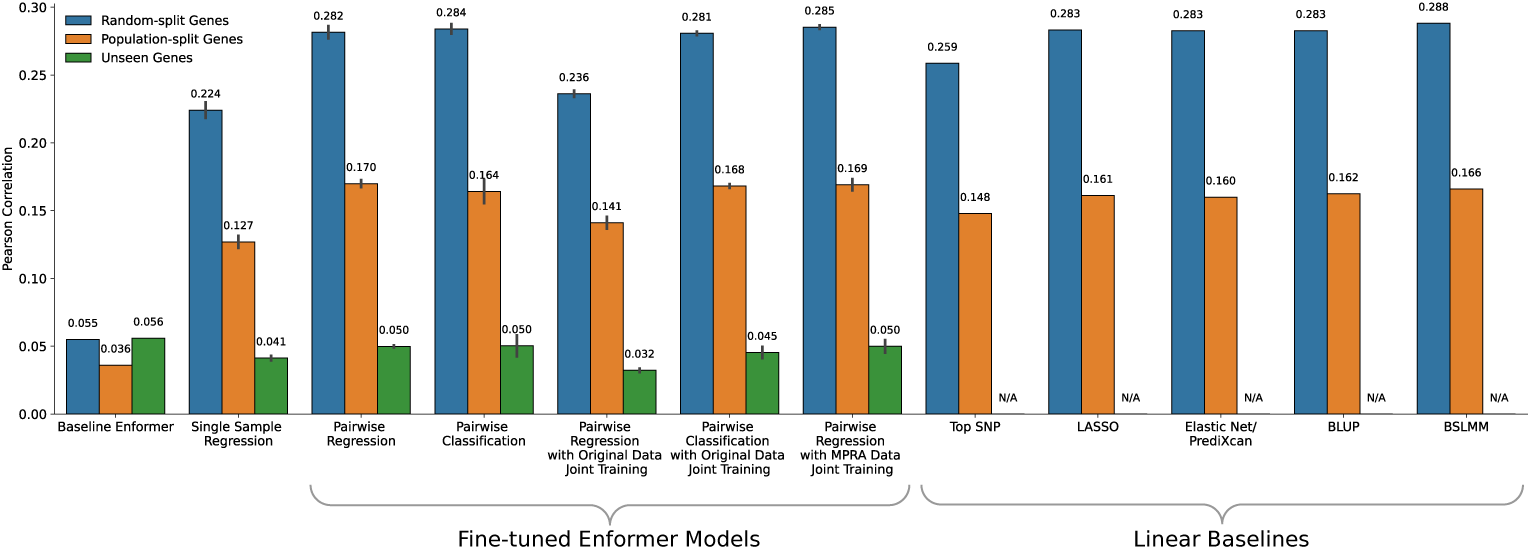
Summary of the performances of all fine-tuned Enformer models and all baseline methods when used to predict individual gene expression levels. Bar heights indicate the mean Pearson correlation across genes. As in Figure 2b, for the fine-tuned models, this mean correlation is averaged across model replicates, and the error bars show the standard deviation among those replicates.

**Fig. S4.**
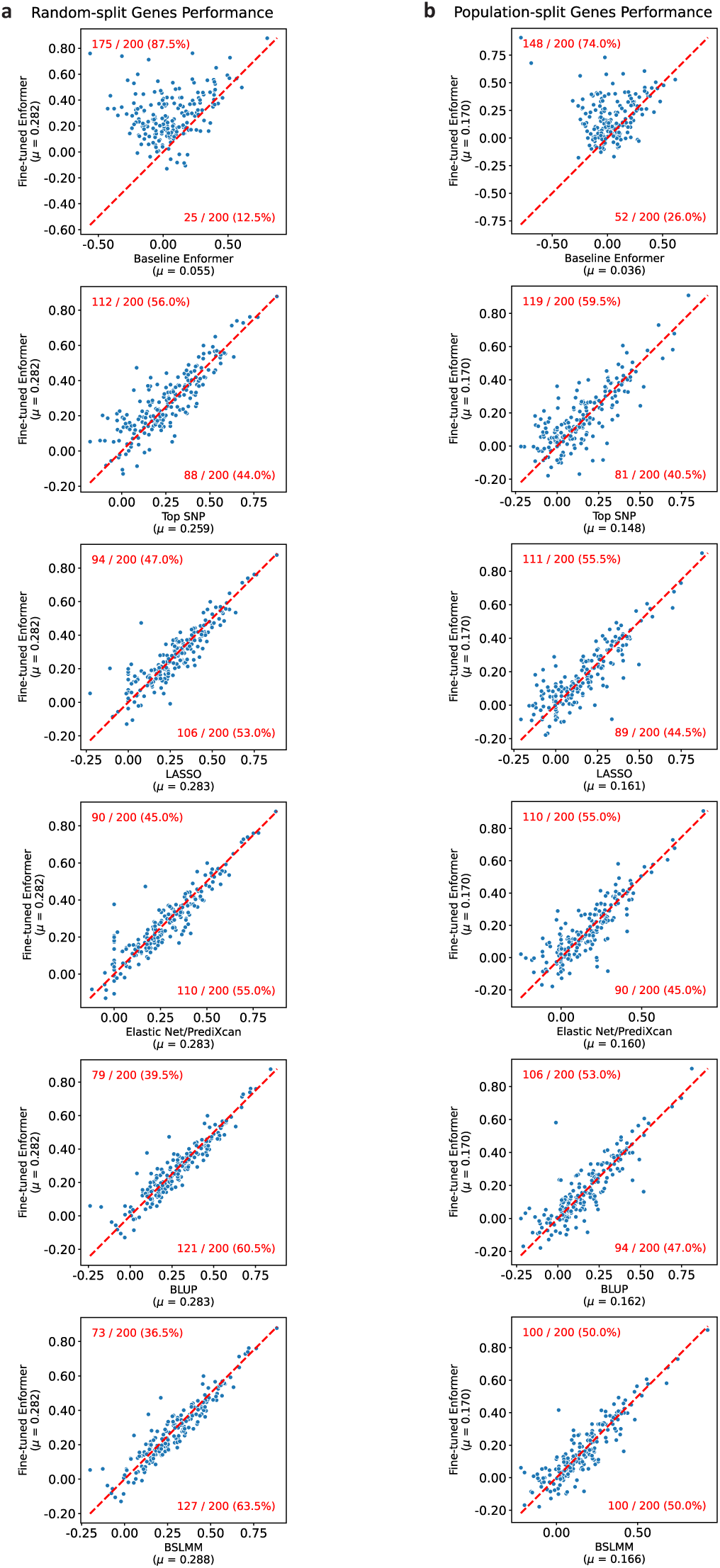
Scatter plots comparing the performance of the fine-tuned Enformer model (pairwise regression) with each baseline method on **(a)** random-split genes and **(b)** population-split genes. Each point represents a gene. The red dashed line indicates *y* = *x*. The number and percentage of genes above and below this line are displayed in red at the top left and bottom right of each plot.

**Fig. S5.**
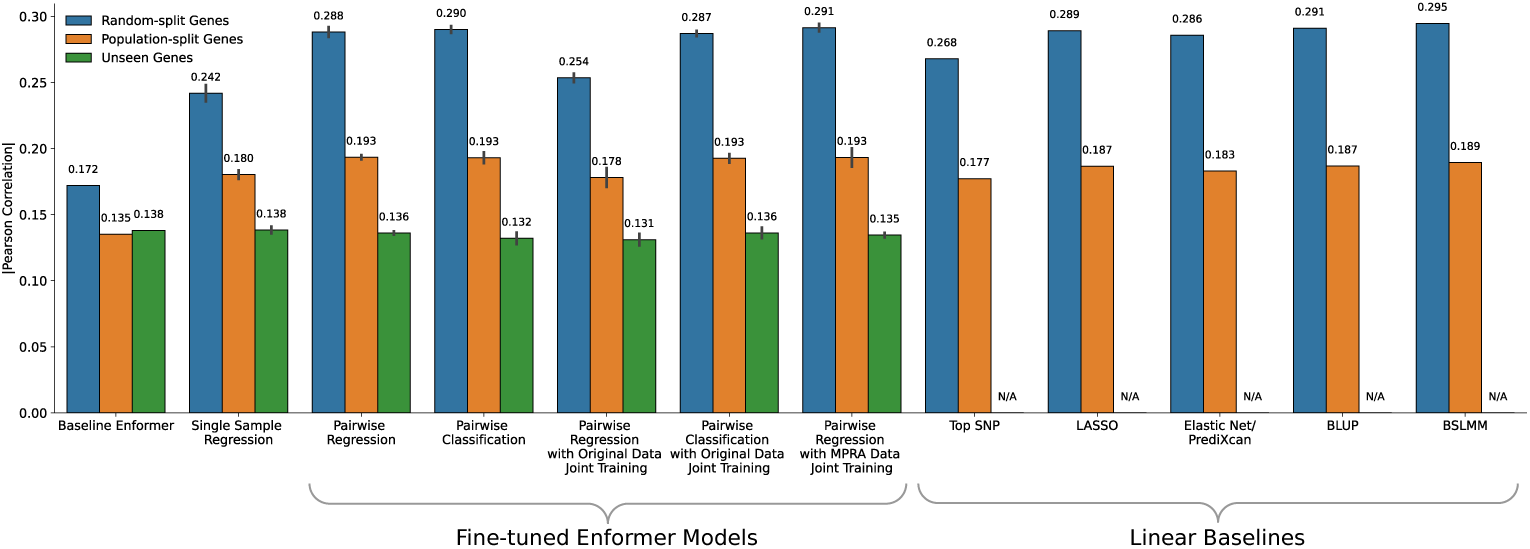
Since models may accurately identify the magnitude of a variant’s effect on expression but not its direction—especially for unseen genes—we compare the absolute values of cross-individual Pearson correlations for all fine-tuned Enformer models and baseline methods. Bar heights indicate the average absolute cross-individual Pearson correlation. As in Fig. 2b, the bar height for fine-tuned models is averaged across three model replicates, with error bars indicating the standard deviation.

**Fig. S6.**
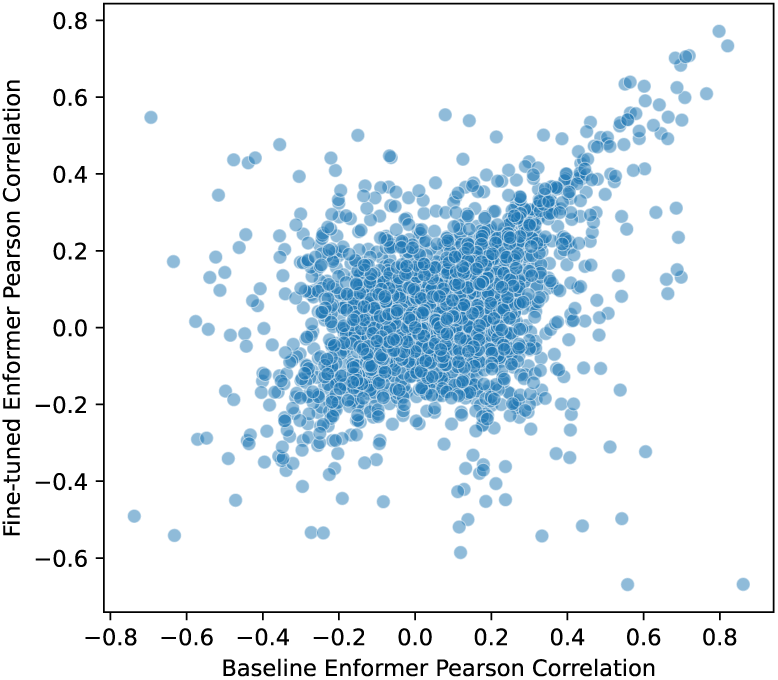
Performance comparison of baseline Enformer and fine-tuned Enformer (pairwise regression) on unseen genes (Spearman correlation = 0.349). Each point in the scatterplot represents an individual gene.

**Fig. S7.**
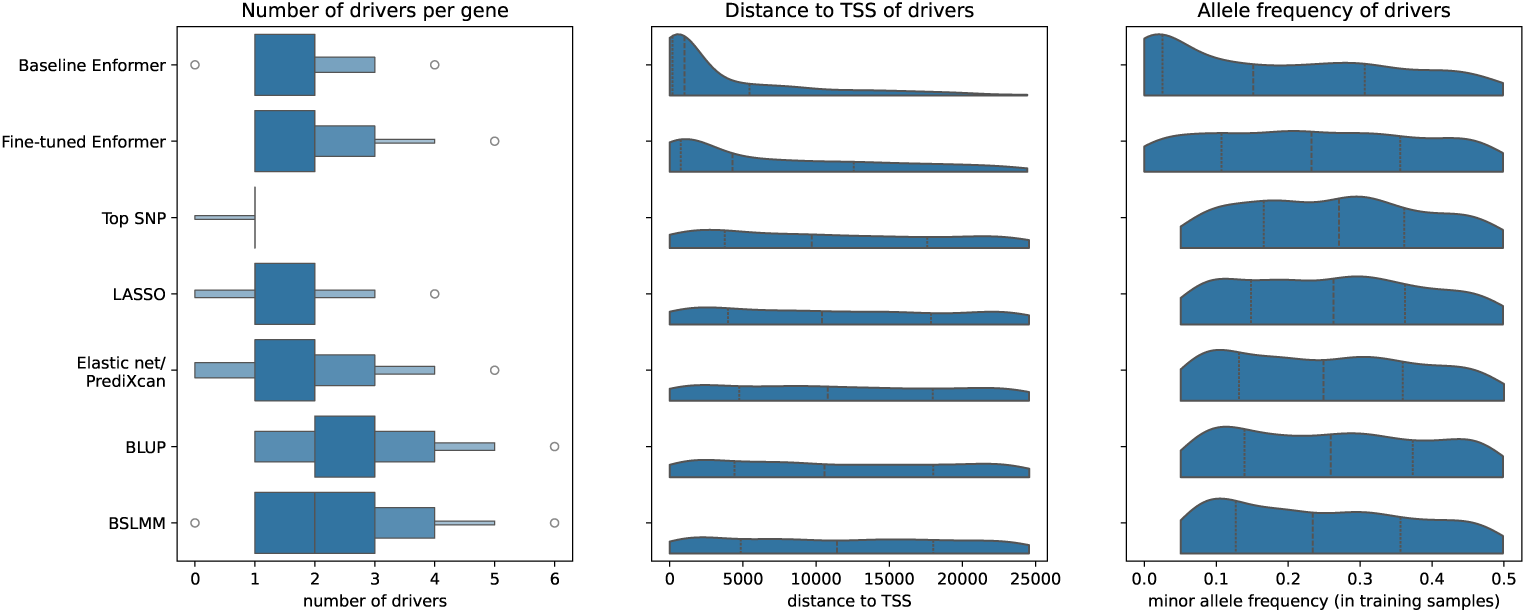
For each model, we identify the variants that drive predictions, termed “drivers,” for both random-split and population-split genes. We analyze (left) the number of drivers per gene, (middle) the unsigned distances of drivers from the transcription start site (TSS) of their corresponding genes, and (right) the minor allele frequencies of drivers, calculated using individuals from the training set. Drivers were identified using the algorithm described in Sasse et al. [12].

## Supplementary Tables

**Table S1.**
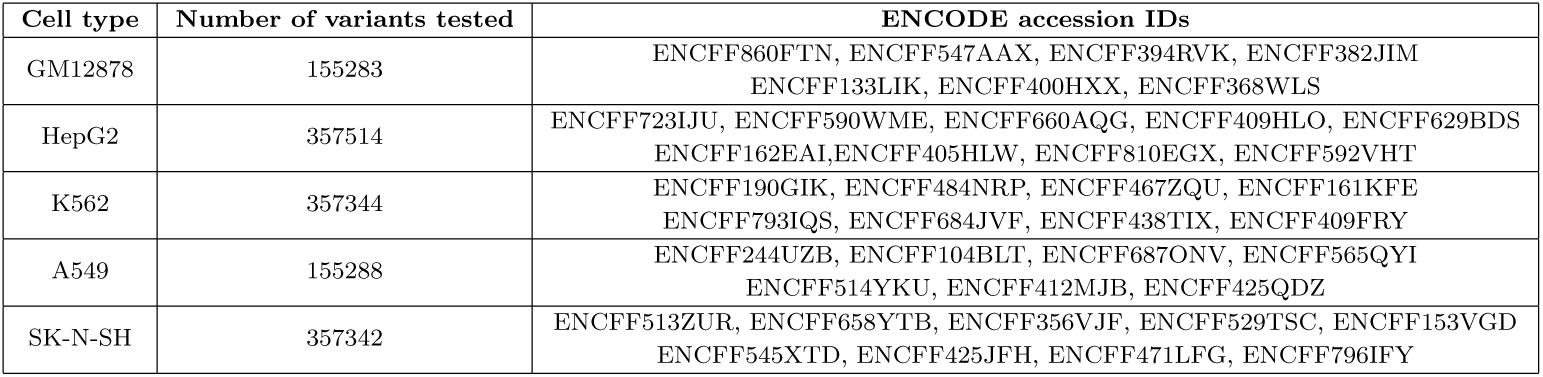
ENCODE accessions IDs used to collate the MPRA data published by Siraj et al. [18].

## References

[1] Gamazon, E.R., Wheeler, H.E., Shah, K.P., Mozaffari, S.V., Aquino-Michaels, K., Carroll, R.J., Eyler, A.E., Denny, J.C., Consortium, G., Nicolae, D.L., et al.: A gene-based association method for mapping traits using reference transcriptome data. Nature genetics 47(9), 1091–1098 (2015)

[2] Gusev, A., Ko, A., Shi, H., Bhatia, G., Chung, W., Penninx, B.W., Jansen, R., De Geus, E.J., Boomsma, D.I., Wright, F.A., et al.: Integrative approaches for large-scale transcriptome-wide association studies. Nature genetics 48(3), 245– 252 (2016)

[3] Zhou, J., Theesfeld, C.L., Yao, K., Chen, K.M., Wong, A.K., Troyanskaya, O.G.: Deep learning sequence-based ab initio prediction of variant effects on expression and disease risk. Nature genetics 50(8), 1171–1179 (2018)

[4] Kelley, D.R., Reshef, Y.A., Bileschi, M., Belanger, D., McLean, C.Y., Snoek, J.: Sequential regulatory activity prediction across chromosomes with convolutional neural networks. Genome research 28(5), 739–750 (2018)

[5] Agarwal, V., Shendure, J.: Predicting mrna abundance directly from genomic sequence using deep convolutional neural networks. Cell reports 31(7), 107663 (2020)

[6] Avsec, Z^̌^., Agarwal, V., Visentin, D., Ledsam, J.R., Grabska-Barwinska, A., Taylor, K.R., Assael, Y., Jumper, J., Kohli, P., Kelley, D.R.: Effective gene expression prediction from sequence by integrating long-range interactions. Nature methods 18(10), 1196–1203 (2021)

[7] Linder, J., Srivastava, D., Yuan, H., Agarwal, V., Kelley, D.R.: Predicting rna-seq coverage from dna sequence as a unifying model of gene regulation. Biorxiv, 2023–08 (2023)

[8] Hernandez, R.D., Uricchio, L.H., Hartman, K., Ye, C., Dahl, A., Zaitlen, N.: Ultrarare variants drive substantial cis heritability of human gene expression. Nature Genetics 51(9), 1349–1355 (2019)

[9] Ferraro, N.M., Strober, B.J., Einson, J., Abell, N.S., Aguet, F., Barbeira, A.N., Brandt, M., Bucan, M., Castel, S.E., Davis, J.R., et al.: Transcriptomic signatures across human tissues identify functional rare genetic variation. Science 369(6509), 5900 (2020)

[10] Schaid, D.J., Chen, W., Larson, N.B.: From genome-wide associations to candidate causal variants by statistical fine-mapping. Nature Reviews Genetics 19(8), 491–504 (2018)

[11] Huang, C., Shuai, R.W., Baokar, P., Chung, R., Rastogi, R., Kathail, P., Ioannidis, N.M.: Personal transcriptome variation is poorly explained by current genomic deep learning models. Nature Genetics 55(12), 2056–2059 (2023)

[12] Sasse, A., Ng, B., Spiro, A.E., Tasaki, S., Bennett, D.A., Gaiteri, C., De Jager, P.L., Chikina, M., Mostafavi, S.: Benchmarking of deep neural networks for predicting personal gene expression from dna sequence highlights shortcomings. Nature Genetics 55(12), 2060–2064 (2023)

[13] Bajwa, A., Rastogi, R., Kathail, P., Shuai, R.W., Ioannidis, N.: Characterizing uncertainty in predictions of genomic sequence-to-activity models. In: Machine Learning in Computational Biology, pp. 279–297 (2024). PMLR

[14] Lappalainen, T., Sammeth, M., Friedländer, M.R., Hoen, P.A., Monlong, J., Rivas, M.A., Gonzalez-Porta, M., Kurbatova, N., Griebel, T., Ferreira, P.G., et al.: Transcriptome and genome sequencing uncovers functional variation in humans. Nature 501(7468), 506–511 (2013)

[15] Mikhaylova, A.V., Thornton, T.A.: Accuracy of gene expression prediction from genotype data with predixcan varies across and within continental populations. Frontiers in genetics 10, 261 (2019)

[16] Keys, K.L., Mak, A.C., White, M.J., Eckalbar, W.L., Dahl, A.W., Mefford, J., Mikhaylova, A.V., Contreras, M.G., Elhawary, J.R., Eng, C., et al.: On the cross-population generalizability of gene expression prediction models. PLoS genetics 16(8), 1008927 (2020)

[17] Taylor, D.J., Chhetri, S.B., Tassia, M.G., Biddanda, A., Yan, S.M., Wojcik, G.L., Battle, A., McCoy, R.C.: Sources of gene expression variation in a globally diverse human cohort. Nature, 1–9 (2024)

[18] Siraj, L., Castro, R.I., Dewey, H., Kales, S., Nguyen, T.T.L., Kanai, M., Berenzy, D., Mouri, K., Wang, Q.S., McCaw, Z.R., et al.: Functional dissection of complex and molecular trait variants at single nucleotide resolution. bioRxiv (2023)

[19] McAfee, J.C., Lee, S., Lee, J., Bell, J.L., Krupa, O., Davis, J., Insigne, K., Bond, M.L., Zhao, N., Boyle, A.P., et al.: Systematic investigation of allelic regulatory activity of schizophrenia-associated common variants. Cell Genomics 3(10) (2023)

[20] Drusinsky, S., Whalen, S., Pollard, K.S.: Deep-learning prediction of gene expression from personal genomes. bioRxiv, 2024–07 (2024)

[21] The 1000 Genomes Project Consortium: A global reference for human genetic variation. Nature 526(7571), 68 (2015)

[22] Zhou, H.J., Li, L., Li, Y., Li, W., Li, J.J.: Pca outperforms popular hidden variable inference methods for molecular qtl mapping. Genome biology 23(1), 210 (2022)

[23] Li, H.: A statistical framework for snp calling, mutation discovery, association mapping and population genetical parameter estimation from sequencing data. Bioinformatics 27(21), 2987–2993 (2011)

[24] Loshchilov, I., Hutter, F.: Decoupled weight decay regularization. In: International Conference on Learning Representations (2019)

[25] McCloskey, M., Cohen, N.J.: Catastrophic interference in connectionist networks: The sequential learning problem. In: Psychology of Learning and Motivation vol. 24, pp. 109–165. Elsevier, New York (1989)

[26] French, R.M.: Catastrophic forgetting in connectionist networks. Trends in cognitive sciences 3(4), 128–135 (1999)

[27] Rusu, A.A., Colmenarejo, S.G., Gulcehre, C., Desjardins, G., Kirkpatrick, J., Pascanu, R., Mnih, V., Kavukcuoglu, K., Hadsell, R.: Policy distillation. arXiv preprint arXiv:1511.06295 (2015)

[28] Parisotto, E., Ba, J.L., Salakhutdinov, R.: Actor-mimic: Deep multitask and transfer reinforcement learning. arXiv preprint arXiv:1511.06342 (2015)

[29] The ENCODE Project Consortium: An integrated encyclopedia of dna elements in the human genome. Nature 489(7414), 57 (2012)

[30] Luo, Y., Hitz, B.C., Gabdank, I., Hilton, J.A., Kagda, M.S., Lam, B., Myers, Z., Sud, P., Jou, J., Lin, K., et al.: New developments on the encyclopedia of dna elements (encode) data portal. Nucleic acids research 48(D1), 882–889 (2020)

[31] Hitz, B.C., Lee, J.-W., Jolanki, O., Kagda, M.S., Graham, K., Sud, P., Gabdank, I., Strattan, J.S., Sloan, C.A., Dreszer, T., et al.: The encode uniform analysis pipelines. Research Square (2023)

[32] Tibshirani, R.: Regression shrinkage and selection via the lasso. Journal of the Royal Statistical Society Series B: Statistical Methodology 58(1), 267–288 (1996)

[33] Zou, H., Hastie, T.: Regularization and variable selection via the elastic net. Journal of the Royal Statistical Society Series B: Statistical Methodology 67(2), 301–320 (2005)

[34] Henderson, C.R.: Best linear unbiased estimation and prediction under a selection model. Biometrics, 423–447 (1975)

[35] Zhou, X., Carbonetto, P., Stephens, M.: Polygenic modeling with bayesian sparse linear mixed models. PLoS genetics 9(2), 1003264 (2013)

[36] Wang, G., Sarkar, A., Carbonetto, P., Stephens, M.: A simple new approach to variable selection in regression, with application to genetic fine mapping. Journal of the Royal Statistical Society Series B: Statistical Methodology 82(5), 1273–1300 (2020)

[37] Broad Institute: Picard Tools. http://broadinstitute.github.io/picard/ (Accessed: 2023/07/13; version 3.0.0.)

